# Interpretable gene networks from single-cell foundation models reveal conserved neurogenic dysfunction in Parkinson’s disease

**DOI:** 10.64898/2026.07.26.740813

**Authors:** Jun Yin, Maya Gosztyla, Birkan Gokbag, Anna Rychkova, Isabel Wilkinson, John St John, Vega Shah, Sarah L. Nickels, Jens C. Schwamborn, Robert T. Fremeau

**Affiliations:** BrainStorm Therapeutics, Inc., San Diego, USA; College of Medicine, The Ohio State University, Columbus, USA; Cosmica Biosciences, San Francisco, USA; NVIDIA Corporation, Santa Clara, USA; University of Luxembourg, Luxembourg, Luxembourg

## Abstract

Interpreting large-scale single-cell transcriptomic data remains a major challenge for understanding disease mechanisms. Recent single-cell foundation models learn rich representations of gene relationships across millions of cells, yet methods for translating these embeddings into biologically interpretable gene networks remain limited. Here we present scGENet, a computational framework that constructs context-specific gene interaction networks from foundation model–derived gene embeddings. By fine-tuning pretrained models on transcriptomic data from human midbrain organoids, scGENet generates transcriptome-scale gene modules that capture biologically meaningful cellular programs. Benchmarking across multiple foundation models demonstrates that networks derived from a fine-tuned scGPT brain model show the highest concordance with curated neuronal pathways, Parkinson’s disease (PD) genetic risk loci, and independent patient-derived transcriptional signatures. Applying this framework to human iPSC-derived PD midbrain organoids reveals transcriptional modules associated with neuronal differentiation, synaptic signaling, and cell-cycle regulation. Single-nucleus RNA sequencing further links these programs to altered cellular composition, including reduced dopaminergic neurons, expansion of radial glia–like progenitors, and a dopaminergic neuron subtype expressing SNCA and VGLUT2. Integration with independent human substantia nigra datasets identifies a conserved neurogenic program disrupted across genetic and idiopathic PD. Together, these results establish a generalizable strategy for extracting interpretable gene networks from single-cell foundation models, enabling systematic discovery of disease-relevant molecular programs across diverse tissues and datasets.

## Introduction

Neurological disorders represent a growing global health burden, yet the development of effective therapies for diseases such as Parkinson’s disease (PD) has been hindered by the limited predictive power of preclinical models. Animal models frequently fail to reproduce key features of human neurodegeneration, contributing to high attrition rates in central nervous system drug development. Human induced pluripotent stem cell (iPSC)-derived brain organoids have emerged as promising systems for modeling neurological disease because they recapitulate aspects of human brain cytoarchitecture, cellular diversity, and developmental trajectories^1,2^. Midbrain organoids in particular enable the generation of dopaminergic neurons and have been widely used to study PD-associated genetic mutations and cellular phenotypes^3,4^.

Despite their promise, extracting mechanistic insight from organoid transcriptomic data remains challenging^5^. Organoid systems generate complex and heterogeneous single-cell datasets that are difficult to interpret using conventional pathway analysis approaches. Traditional enrichment analyses rely on predefined gene sets that are often redundant and lack cell-type specificity, limiting their ability to reveal context-dependent molecular programs^6,7^. Similarly, conventional gene co-expression methods such as weighted gene co-expression network analysis (WGCNA)^8^ are constrained by sparse gene detection in single-cell datasets and typically capture only a fraction of the transcriptome.

Recent advances in single-cell foundation models, including scGPT^9^, Geneformer^10^, and scFoundation^11^, provide new opportunities to address these limitations. Trained on tens of millions of single-cell transcriptomes, these models learn high-dimensional representations of gene relationships that encode latent biological structure^9–12^. While foundation models have been successfully applied to tasks such as cell-type annotation, batch correction, and perturbation prediction, relatively few studies have explored how gene embeddings derived from these models can be used to construct transcriptome-scale gene interaction networks^13,14^.

Here we introduce scGENet, a computational framework that leverages gene embeddings from single-cell foundation models to construct context-specific gene networks directly from experimental data. By fine-tuning pretrained models on transcriptomic data from human midbrain organoids, scGENet generates biologically interpretable gene modules that capture disease-relevant transcriptional programs. We systematically benchmark multiple foundation models and demonstrate that fine-tuned scGPT-based networks outperform conventional co-expression approaches in capturing neuronal biological processes, PD genetic risk signals, and patient-derived transcriptional signatures.

Applying this framework to human iPSC-derived midbrain organoids carrying PD-associated mutations, we identify transcriptional modules associated with neuronal differentiation, synaptic signaling, and cell-cycle regulation. Integration with single-nucleus RNA sequencing and independent human brain datasets reveals a conserved transcriptional program related to neuronal maturation that is disrupted across genetic and idiopathic PD. Together, these findings establish foundation model–derived gene networks as a scalable approach for interpreting complex transcriptomic datasets and linking experimental disease models with molecular programs observed in human neurodegenerative disease.

## Results

### scGENet constructs gene interaction networks from single-cell foundation model embeddings

Interpreting large-scale single-cell transcriptomic data from complex biological systems such as human brain organoids remains challenging. To address this limitation, we developed scGENet, a computational framework that constructs gene interaction networks from embeddings generated by single-cell foundation models.

To establish a disease model, midbrain organoids were generated from four control lines, one familial PD line harboring an LRRK2 mutation^15^, and seven familial PD lines carrying GBA1 mutations^16^ (Fig. 1A). These organoids recapitulate key neuronal and glial populations of the developing human midbrain and provide a scalable platform for transcriptomic analysis.

**Figure 1.**
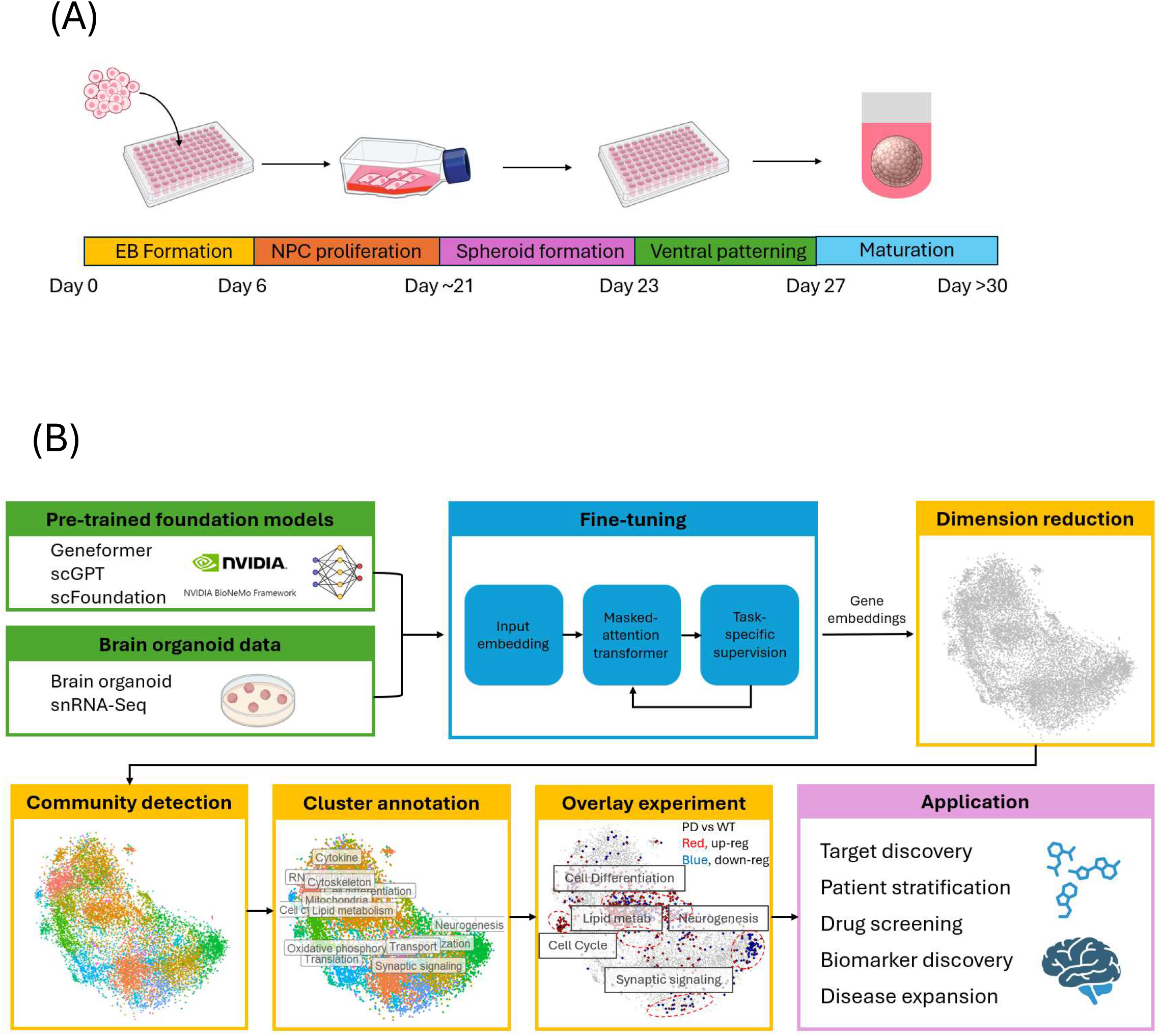
scGENet constructs gene interaction networks from single- cell foundation model embeddings. **(A)** Schematic of human iPSC-derived midbrain organoid generation from control and Parkinson’s disease (PD) patient lines used for transcriptomic profiling. **(B)** Overview of the scGENet framework, in which gene embeddings derived from fine-tuned single-cell foundation models are used to construct transcriptome-scale gene interaction networks and identify biologically meaningful gene modules. The gene network was constructed using fine-tuned scGPT brain model. The gene expression changes in overlay experiment were from PD vs WT midbrain organoid bulk RNA-Seq.

scGENet constructs context-specific gene networks from gene embeddings derived from single-cell foundation models trained on tens of millions of transcriptomes^9–11^. Unlike conventional co-expression approaches that infer gene relationships directly from expression correlations^8^, scGENet leverages latent gene representations learned across large single-cell datasets, enabling transcriptome-scale inference of gene relationships even for genes with sparse expression.

The framework consists of three steps (Fig. 1B). First, pretrained foundation models are fine-tuned using transcriptomic data from the biological system of interest, enabling gene embeddings to capture context-specific transcriptional relationships. Second, these embeddings are used to construct a transcriptome-wide gene interaction network through nearest-neighbor graph construction followed by community detection to identify gene modules. Third, the resulting modules are evaluated through multiple biological benchmarks, including enrichment for curated pathway databases, concordance with disease-associated genetic loci, agreement with independent transcriptomic datasets, and network topology metrics.

Applying scGENet to single-nucleus transcriptomic data from control and PD midbrain organoids produced gene interaction networks capturing transcriptional programs associated with neuronal differentiation, synaptic signaling, metabolism, and cell cycle regulation (Fig. 1B). We next evaluated the performance of different foundation models for constructing biologically informative gene networks.

### Benchmarking identifies scGPT as the most informative representation for gene network construction

To assess the ability of foundation models to generate biologically meaningful gene networks, we benchmarked networks derived from several pretrained single-cell models, including scGPT^9^, Geneformer^10^, and scFoundation^11^. Networks were generated both with and without fine-tuning on midbrain organoid single-nucleus sequencing data. For comparison, the conventional single-cell co-expression method hdWGCNA^8^ was also included as a baseline approach for network construction.

Gene embeddings from each model were used to construct gene networks using the scGENet workflow, and resulting gene modules were evaluated using multiple orthogonal criteria (Fig 2. A-D). These included enrichment for curated Gene Ontology (GO) biological processes, concordance with PD GWAS loci^17^, and overlap with differentially expressed genes (DEGs) derived from midbrain organoid bulk RNA-seq datasets.

**Figure 2.**
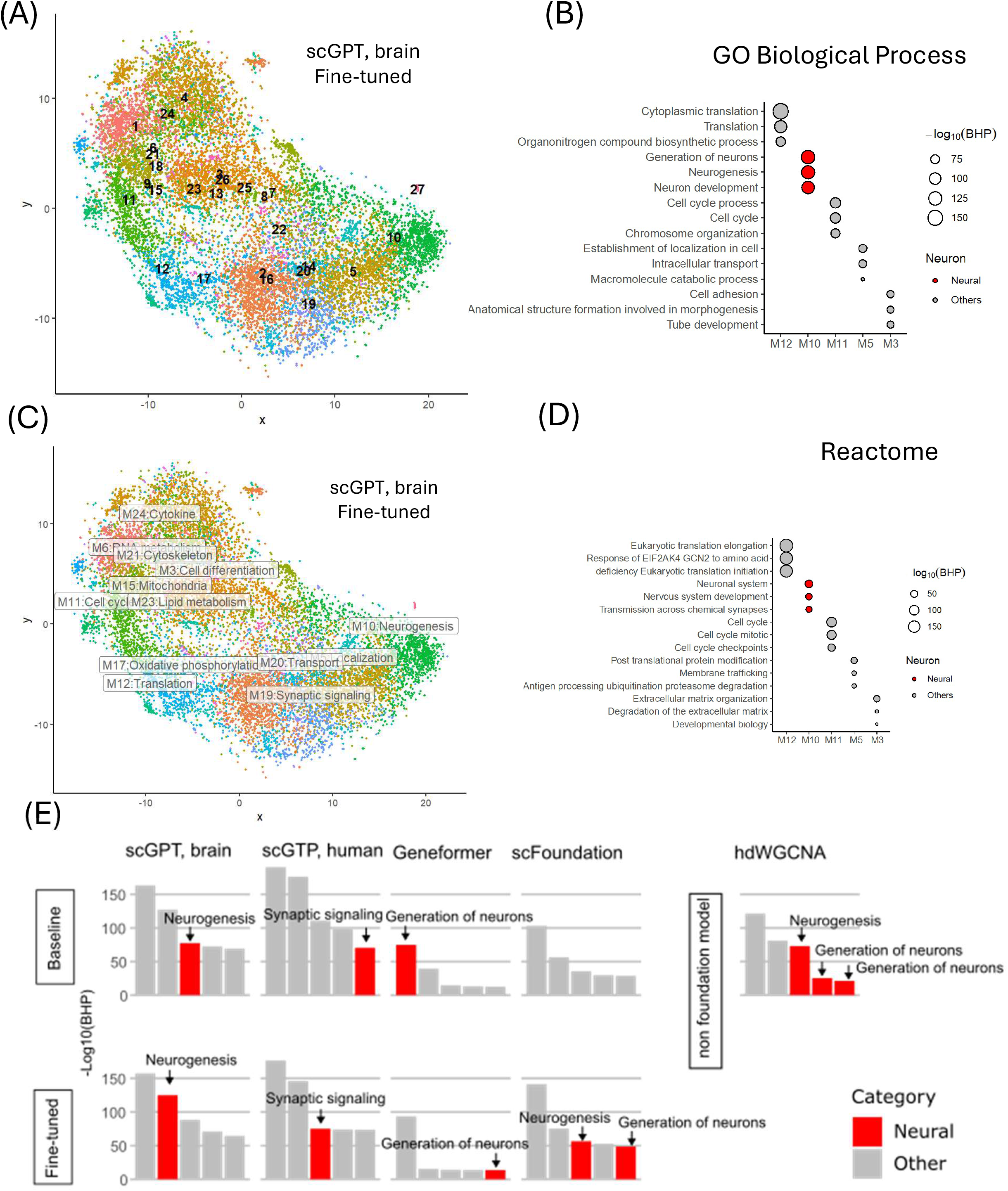
Gene module identification and benchmarking of foundation model–derived gene networks. (A) Visualization of gene modules derived from foundation model embeddings using t-SNE dimensionality reduction. (B) Gene Ontology (GO) biological process enrichment analysis for the top five gene modules ranked by enrichment score. (C) Functional annotation of gene modules based on enriched biological processes. (D) Reactome pathway enrichment analysis of the top five gene modules. (E) Benchmarking of gene module enrichment across networks generated using pre-trained and fine- tuned foundation models and the hdWGCNA method. Modules associated with neuronal biological processes are highlighted in red. Bar plots show the top five modules ranked by GO enrichment score for each network.

Across these benchmarks, gene networks derived from the fine-tuned scGPT brain model consistently showed the highest concordance with biological benchmarks (Fig 2E, Supplementary Fig. 1-3). Fine-tuning the scGPT foundation model further improved network agreement with established biological knowledge. In particular, modules derived from the fine-tuned scGPT networks demonstrated higher enrichment for neuron-associated GO terms compared with other models and conventional co-expression networks (Fig. 2E). In addition, these modules showed stronger enrichment for PD-associated GWAS signals (P < 1 × 10⁻¹⁵) (Supplementary Fig. 1A,B) and greater concordance with transcriptional signatures derived from PD organoid bulk RNA-seq datasets (Supplementary Fig. 1C).

In contrast, the hdWGCNA approach requires genes to be expressed in more than 5% of cells, substantially limiting the number of genes included in the network. As a result, hdWGCNA incorporated only 3,951 genes into five modules, whereas foundation model–derived networks captured more than 16,000 genes organized into over 20 modules. These results indicate that foundation model embeddings enable more comprehensive and biologically informative gene network construction from single-cell datasets.

To further evaluate network quality, we analyzed structural properties of the networks using topology metrics. Gene networks constructed using scGPT models, with or without fine-tuning, exhibited higher small-worldness and Gini coefficients compared with networks generated by other foundation models. These properties are consistent with the modular and hierarchically organized architecture characteristic of biological networks. However, fine-tuning did not appear to substantially alter the overall network structural properties (Supplementary Fig. 3).

Based on these results, the fine-tuned scGPT brain network was selected as the reference gene network for subsequent analyses and served as the reference framework for mapping disease-associated transcriptional changes in PD organoids (Fig. 2 A-D).

### Gene network analysis identifies dysregulated neurogenesis programs in PD organoids

Having established a foundation model–derived gene network, we next applied this framework to identify disease-associated transcriptional programs in PD organoids.

To evaluate transcriptomic reproducibility, bulk RNA sequencing was performed on day-30 midbrain organoids derived from three GBA1 PD patients and three control donors across six independent experimental batches. Transcriptomic profiles showed high reproducibility, with pairwise correlation coefficients exceeding 0.92 across samples (Fig. 3A). Principal variance component analysis indicated that gene expression variance was primarily driven by donor identity, disease status, and sex, with minimal contribution from batch effects (Fig. 3B).

**Figure 3.**
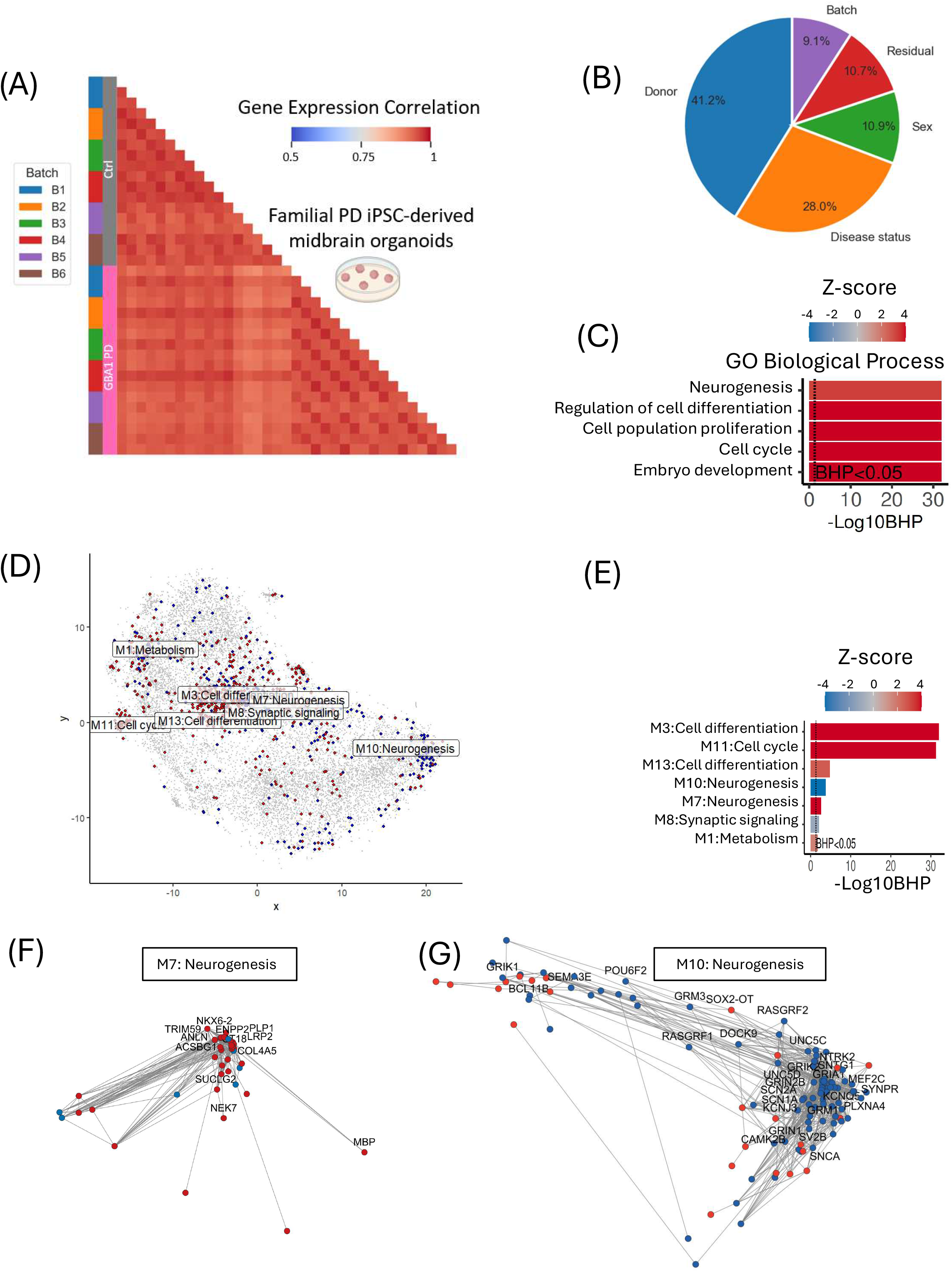
Foundation model–derived network analysis reveals transcriptional modules altered in PD midbrain organoids. (A) Pearson correlation of bulk RNA-seq samples from three GBA1 PD patients and three control donors across six experimental batches. (B) Principal variance component analysis (PVCA) showing contributions of donor, disease status, and other factors to gene expression variance. (C) Gene Ontology enrichment analysis of differentially expressed genes (DEGs) between PD and control midbrain organoids. Positive Z-scores indicate enrichment of upregulated genes and negative Z-scores indicate enrichment of downregulated genes. (D) Visualization of DEGs mapped onto the foundation model–derived gene network. Upregulated genes are shown in red and downregulated genes in blue. Gene modules significantly enriched with DEGs (BH-adjusted P < 0.05) are labeled. (E) Enrichment of PD-associated DEGs across gene network modules. (F–G) Differentially expressed genes within neurogenesis-associated modules M10 and M7 comparing GBA1 PD and control organoids. The positions of the genes were according to the t-SNE coordinates in Fig. 3D.

Differential expression analysis between PD and control organoids revealed widespread transcriptional changes. Conventional pathway enrichment analysis using Gene Ontology^6^, Reactome^7^, WikiPathways^18^, and PanglaoDB^19^ identified enrichment for processes associated with neurogenesis, cell differentiation, and cell cycle regulation (Fig. 3C and Supplementary Fig. 4). However, these predefined gene sets often overlap substantially and do not capture context-specific gene relationships.

Projecting differentially expressed genes onto the foundation model–derived gene network resolved these signals into distinct transcriptional modules with coherent biological themes (Fig. 3D–E). These included modules associated with cell cycle regulation, neuronal differentiation, synaptic signaling, neurogenesis, and metabolism.

Notably, genes annotated under the GO neurogenesis term were distributed across multiple network modules, including M3 (cell differentiation), M7, and M10, each showing distinct expression patterns (Supplementary Fig. 5). The network thus separated transcriptional programs that are conflated in conventional enrichment analyses.

Module M7 appeared to represent a progenitor or glial lineage program, containing progenitor-associated transcription factors and glial markers such as LRP2, ST18, and NKX6-2, along with oligodendrocyte-associated genes MBP and PLP1 (Fig. 3F). In contrast, module M10 corresponded to a transcriptional program associated with neuronal maturation and synaptic function, including genes involved in neurotransmission (GRIA1, SCN1A, KCNQ5), synaptic vesicle biology (SNCA, SV2B, SYNPR), and neuronal differentiation (BCL11B, MEF2C, POU6F2) (Fig. 3G).

A separate module (M11) was enriched for canonical cell-cycle regulators, including CDK1, CCNB1, FOXM1, and MKI67, and was predominantly upregulated in PD organoids. These findings demonstrate that foundation model–derived networks resolve PD-associated transcriptional changes into biologically interpretable gene modules.

To determine whether these transcriptional differences reflect changes in cellular composition, we next performed single-nucleus RNA sequencing.

### Single-nucleus RNA sequencing reveals cell-type shifts in PD organoids

Single-nucleus RNA sequencing of PD and control midbrain organoids revealed major neuronal and glial populations, including dopaminergic neurons, GABAergic neurons, radial glia, and neural progenitor cells (Fig. 4A–B and Supplementary Fig. 6).

**Figure 4.**
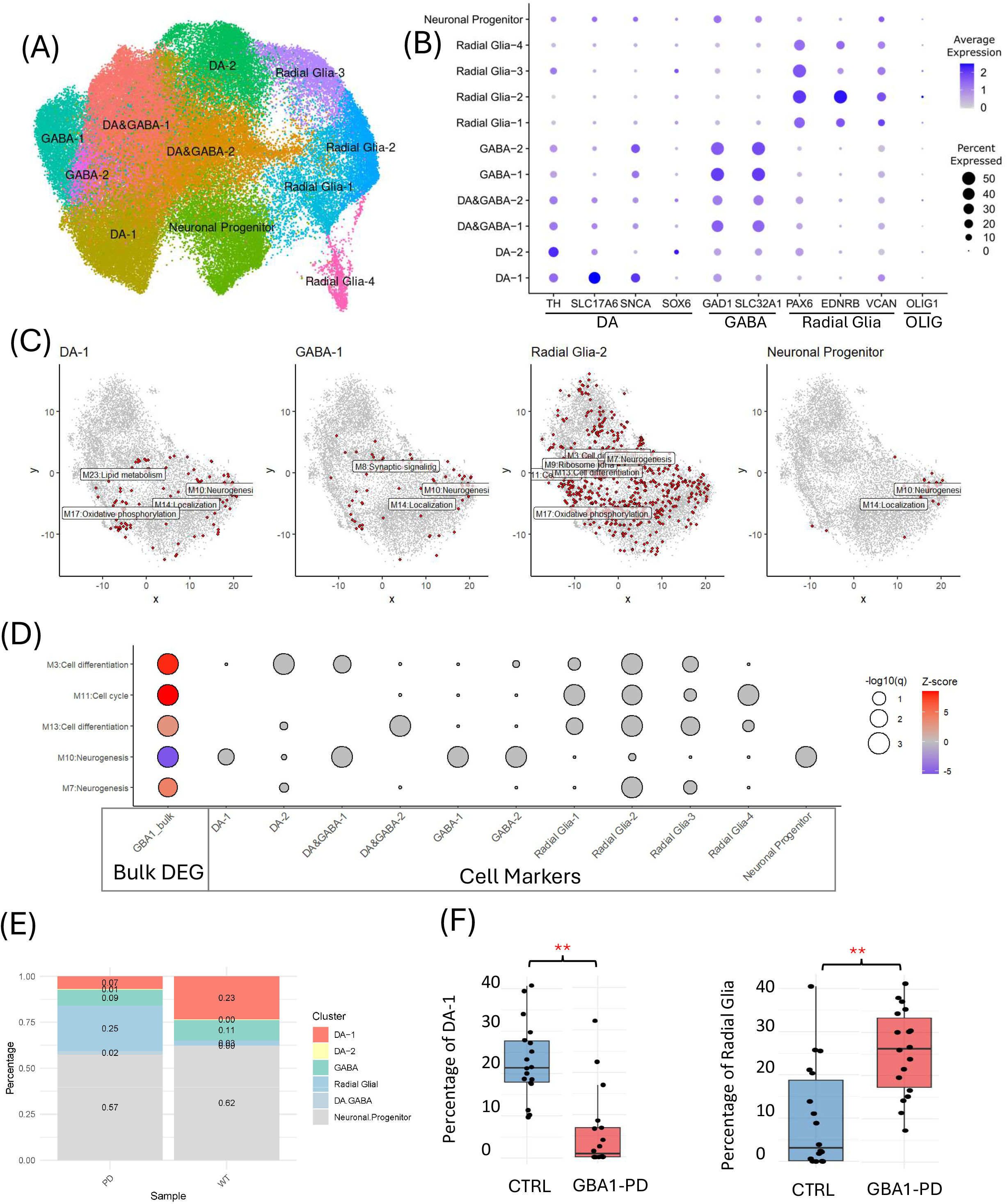
Single-nucleus RNA sequencing reveals cell-type shifts in PD midbrain organoids. (A) UMAP visualization of cell clusters identified from snRNA-seq analysis of PD and control midbrain organoids. (B) Expression of representative cell-type marker genes across identified clusters. DA (Dopaminergic neurons), GABA(GABAergic neurons), and OLIG (Oligodendrocytes) (C) Mapping of cell-type marker genes onto the foundation model–derived gene network. Only modules significantly enriched for cell markers (BH-adjusted P < 0.05) are labeled. (D) Comparison of gene module enrichment scores for cell-type markers and bulk RNA-seq DEGs. (E) Estimated cell-type composition in bulk RNA-seq samples inferred by cell-type deconvolution using snRNA-seq data as reference. (F) Proportions of dopaminergic neurons and radial glial cells in PD and control organoids. **P < 0.01.

Among dopaminergic populations, four clusters were identified: two dopaminergic neuron clusters (DA- 1 and DA-2) and two hybrid dopaminergic/GABAergic (DA&GABA) clusters. The DA-1 cluster expressed TH, VGLUT2, and SNCA, consistent with a glutamate-co-releasing dopaminergic neuron subtype^20^.

Pseudotime trajectory analysis revealed differentiation trajectories from neural progenitors toward mature neuronal lineages and a distinct branch corresponding to radial glial cells (Supplementary Fig. 7).

Mapping cell-type markers onto the gene network showed that dopaminergic neuron and neuronal progenitor markers were enriched in the M10 neurogenesis module, which was downregulated in PD organoids (Fig. 4C–D). In contrast, radial glial markers mapped predominantly to the M7 neurogenesis and M11 cell-cycle modules, which were upregulated in PD organoids.

To quantify changes in cell composition in the midbrain organoid bulk RNA-Seq, we performed FARDEEP-based cell-type deconvolution using single-cell data as a reference^21^. This analysis revealed a significant reduction in dopaminergic neuron abundance and a corresponding increase in radial glial populations in PD organoids compared with controls (Fig. 4E–F).

These findings suggest that transcriptional changes observed in PD organoids arise in part from altered differentiation dynamics, including impaired dopaminergic neuron maturation and expansion of proliferative glial progenitors.

### Foundation model–derived gene networks reveal conserved neurogenic dysfunction across PD models

We next asked whether the transcriptional programs identified in PD organoids are conserved across independent PD models and human brain datasets.

Differential expression profiles were compared across GBA1 PD organoids, LRRK2 PD organoids, and two independent datasets of idiopathic PD substantia nigra tissue, including the Smajic et al. dataset^22^ and the Aligning Science Across Parkinson’s (ASAP) dataset^23^.

Across these datasets, we observed consistent downregulation of genes associated with neurogenesis, synaptic signaling, and neuronal differentiation in dopaminergic neurons (Fig. 5A and Supplementary Fig. 9). The M10 neurogenesis module was significantly enriched with differentially expressed genes in all four datasets (BH-adjusted P < 0.05).

**Figure 5.**
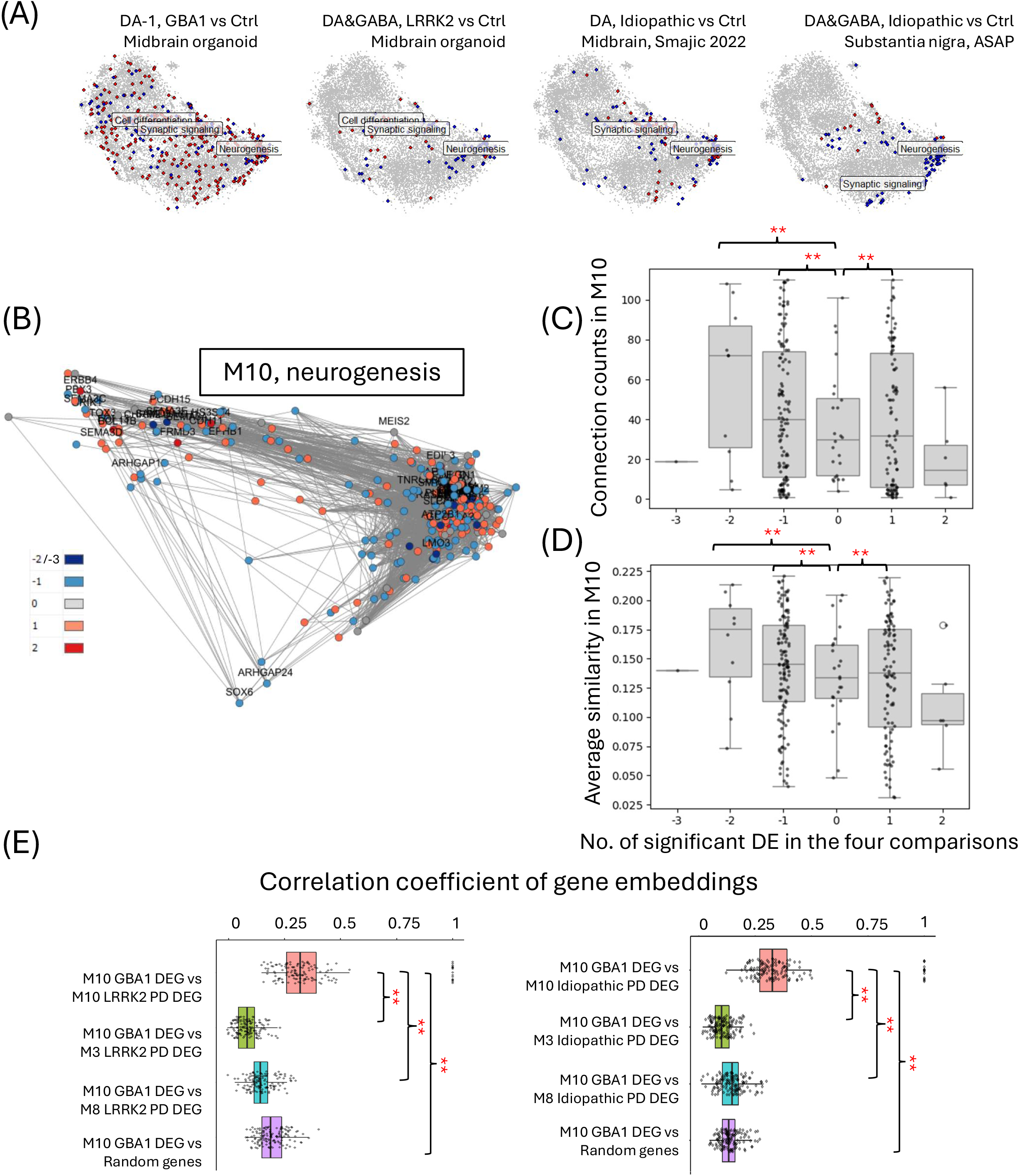
Cross-dataset gene network analysis identifies conserved transcriptional programs across PD models. (A) Differential gene expression in dopaminergic neurons comparing PD and control samples across GBA1 organoids, LRRK2 organoids, and two idiopathic PD brain datasets (Smajic et al. and ASAP). Red indicates upregulated genes and blue indicates downregulated genes. (B) Network visualization of genes differentially expressed in dopaminergic neurons across the four datasets. Colors indicate the number and direction of differential expression events across datasets. -2 or -3 as dark blue, down-regulated in two or three of the four above comparisons. -1, as light blue, down-regulated in one of the four above comparisons. 1, as light red, up-regulated in one of the four above comparisons. 2, as dark red, up-regulated in two of the four above comparisons. 0, as gray, opposite direction in two of the four above comparisons. (C–D) Number of network connections (cosine similarity > 0.25) and average cosine similarity for genes grouped by differential expression category. **P < 0.01. (E) Spearman correlation of DEGs within the neurogenesis module (M10) from GBA1 organoids compared with DEGs in dopaminergic neurons from LRRK2 PD and idiopathic PD brain datasets. **P < 0.01.

Although no individual genes were differentially expressed across all datasets simultaneously, genes downregulated in at least two datasets exhibited significantly higher network connectivity and cosine similarity compared with randomly selected genes (Wilcoxon rank-sum P < 0.01) (Fig. 5B–D). These genes included regulators of neuronal connectivity and differentiation such as TENM2, SEMA3E, CHL1, SAMD5, PDE4D, and CADPS, as well as the transcription factor LMO3, which was downregulated in three datasets.

Correlation analysis further demonstrated that differentially expressed genes within the M10 neurogenesis module showed significantly higher cross-dataset correlation compared with genes from other modules or randomly selected genes (Supplementary Fig. 10).

Together these results demonstrate that foundation model–derived gene networks capture conserved transcriptional programs associated with Parkinson’s disease across organoid models and human brain tissue, highlighting disrupted neurogenesis as a shared molecular feature of PD pathophysiology.

### Results summary

Foundation model–derived gene networks generated using scGENet provide a scalable framework for interpreting complex single-cell transcriptomic datasets and resolving disease-relevant transcriptional programs that are not readily captured by conventional pathway or co-expression analyses. By integrating large-scale foundation model representations with organoid transcriptomics and independent human brain datasets, this approach reveals conserved molecular programs underlying Parkinson’s disease and provides a general strategy for linking experimental disease models with human neurodegenerative biology.

## Discussion

In this study, we present scGENet, a computational framework that leverages single-cell foundation models to construct biologically interpretable gene interaction networks from transcriptomic data. By combining pretrained representations learned from millions of single cells with context-specific fine-tuning on organoid datasets, scGENet enables transcriptome-scale reconstruction of gene networks that capture disease-relevant molecular programs. Applying this approach to human midbrain organoids modeling Parkinson’s disease (PD), we demonstrate that foundation model–derived gene networks can resolve transcriptional modules associated with neuronal differentiation, synaptic signaling, and cell cycle regulation, and identify conserved neurogenic programs disrupted across genetic and idiopathic forms of PD.

Recent advances in large-scale foundation models for single-cell biology have enabled the learning of rich representations of gene relationships across diverse tissues and experimental conditions^9–12^. However, most applications of these models have focused on tasks such as cell-type annotation, batch correction, or perturbation prediction. Methods for translating foundation model embeddings into interpretable gene interaction networks remain relatively unexplored. Our results demonstrate that embeddings derived from these models can be used to construct biologically meaningful networks that capture transcriptional programs beyond those accessible through conventional pathway enrichment or co-expression analysis. In particular, we show that networks derived from a fine-tuned scGPT brain model exhibit strong concordance with curated biological processes, PD genetic risk loci, and independent patient-derived transcriptomic datasets.

Compared with traditional co-expression approaches, foundation model–derived networks provide several advantages. First, because the embeddings are trained on tens of millions of cells, they encode transcriptome-wide gene relationships that are not limited by sparsity in individual single-cell datasets. Second, fine-tuning allows the embeddings to capture context-specific transcriptional relationships, enabling network construction tailored to the tissue and disease state under investigation. Third, the resulting networks capture substantially larger fractions of the transcriptome, facilitating identification of disease-associated modules that may be missed by conventional approaches that rely on expression thresholds.

Recent single-cell foundation models learn gene representations from millions of transcriptomes spanning diverse tissues, developmental stages, and experimental contexts. These models therefore capture latent biological structure reflecting conserved relationships between genes and cellular programs. In this study, we demonstrate that these learned representations can serve as informative priors for gene network reconstruction. By fine-tuning foundation models on midbrain organoid datasets, scGENet adapts these global representations to a specific disease context while retaining information learned from large-scale single-cell data. This approach enables identification of transcriptional modules that extend beyond direct expression correlations observed in individual datasets.

Applying this framework to PD midbrain organoids revealed transcriptional programs associated with impaired neuronal differentiation and altered glial dynamics. In particular, network analysis resolved the broadly defined Gene Ontology neurogenesis category into distinct modules with opposing expression patterns, highlighting the ability of foundation model–derived networks to disentangle overlapping biological pathways. Integration with single-nucleus RNA sequencing further linked these transcriptional programs to cellular composition changes, including reduced dopaminergic neuron abundance and expansion of radial glia–like progenitors. Among dopaminergic populations, we observed a neuronal subtype expressing TH, VGLUT2, and SNCA, consistent with glutamate-co-releasing dopaminergic neurons that have been implicated in synaptic vulnerability and metabolic stress in neurodegenerative contexts^24^.

Importantly, the transcriptional modules identified in PD organoids were conserved across independent experimental systems. Cross-dataset analyses revealed consistent downregulation of genes associated with neuronal differentiation and synaptic signaling across GBA1 and LRRK2 organoid models as well as idiopathic PD midbrain tissue. The convergence of these datasets on a shared neurogenesis module suggests that disrupted neuronal maturation and connectivity may represent a common molecular feature of PD pathophysiology. These findings illustrate how integrating organoid models with foundation model–derived gene networks can bridge experimental systems and human disease biology.

More broadly, our results highlight the potential of combining AI-based representation learning with human organoid models to accelerate disease mechanism discovery. Organoid systems capture key aspects of human tissue architecture and developmental trajectories, but interpreting their complex transcriptomic landscapes remains challenging. By providing a scalable approach for extracting biologically interpretable gene networks from organoid datasets, scGENet offers a general framework for linking experimental disease models to transcriptional programs observed in human tissues.

An important distinction in complex diseases such as Parkinson’s disease is the difference between genes that contribute to disease susceptibility and transcriptional programs that reflect disease progression or cellular response. Genome-wide association studies identify loci that influence disease risk, but these genes represent only a subset of the broader molecular pathways perturbed during disease development. In contrast, the transcriptional modules identified by scGENet capture coordinated gene expression programs that reflect the cellular state of dopaminergic neurons and their supporting cell populations. Notably, while several PD-associated genes were enriched within these modules, the network structure also revealed additional genes that were not individually associated with disease risk but participate in shared biological programs. This highlights the ability of foundation model–derived gene networks to connect genetic susceptibility with downstream molecular processes that shape disease progression.

Several limitations should be considered. First, while foundation model embeddings capture statistical relationships between genes, they do not directly encode causal regulatory interactions. Integrating embedding-derived networks with perturbational datasets or curated molecular interaction networks may further improve mechanistic interpretation. Second, although we evaluated several widely used foundation models, the rapidly evolving landscape of single-cell foundation models means that future models trained on larger and more diverse datasets may yield further improvements. Finally, our analyses focused on transcriptional programs associated with PD organoids at specific developmental stages; extending this framework to longitudinal organoid maturation or perturbational experiments may provide additional insights into disease progression.

Recent graph-based transformer architectures have shown promise for mechanistic reasoning over curated molecular interaction networks^25,26^. However, the objective of this study was to learn transferable representations of disease-associated transcriptional programs directly from single-cell transcriptomic state, without relying on predefined network structures. By fine-tuning foundation models on midbrain organoid datasets, scGENet captures gene expression programs associated with Parkinson’s disease that generalize across experimental systems and patient cohorts. Future work may integrate foundation model–derived embeddings with interactome-level graph reasoning to further connect transcriptional disease states with causal regulatory circuits and therapeutic hypotheses.

Despite these limitations, our study establishes foundation model–derived gene networks as a powerful approach for interpreting complex transcriptomic datasets. By combining large-scale pretrained representations with experimental disease models, this framework enables systematic discovery of transcriptional programs that connect cellular phenotypes with human disease biology. Because scGENet operates directly on gene embeddings derived from foundation models, it is broadly applicable to diverse tissues, organoid systems, and disease contexts.

Together, these results demonstrate how foundation model–based network analysis can bridge artificial intelligence and experimental biology, providing a scalable strategy for uncovering molecular mechanisms in complex diseases such as Parkinson’s disease. As large-scale single-cell datasets and foundation models continue to expand, approaches that integrate representation learning with experimental systems will become increasingly important for translating complex transcriptomic data into biological insight.

## Supplementary Figure Legends

**Supplementary Figure 1.**
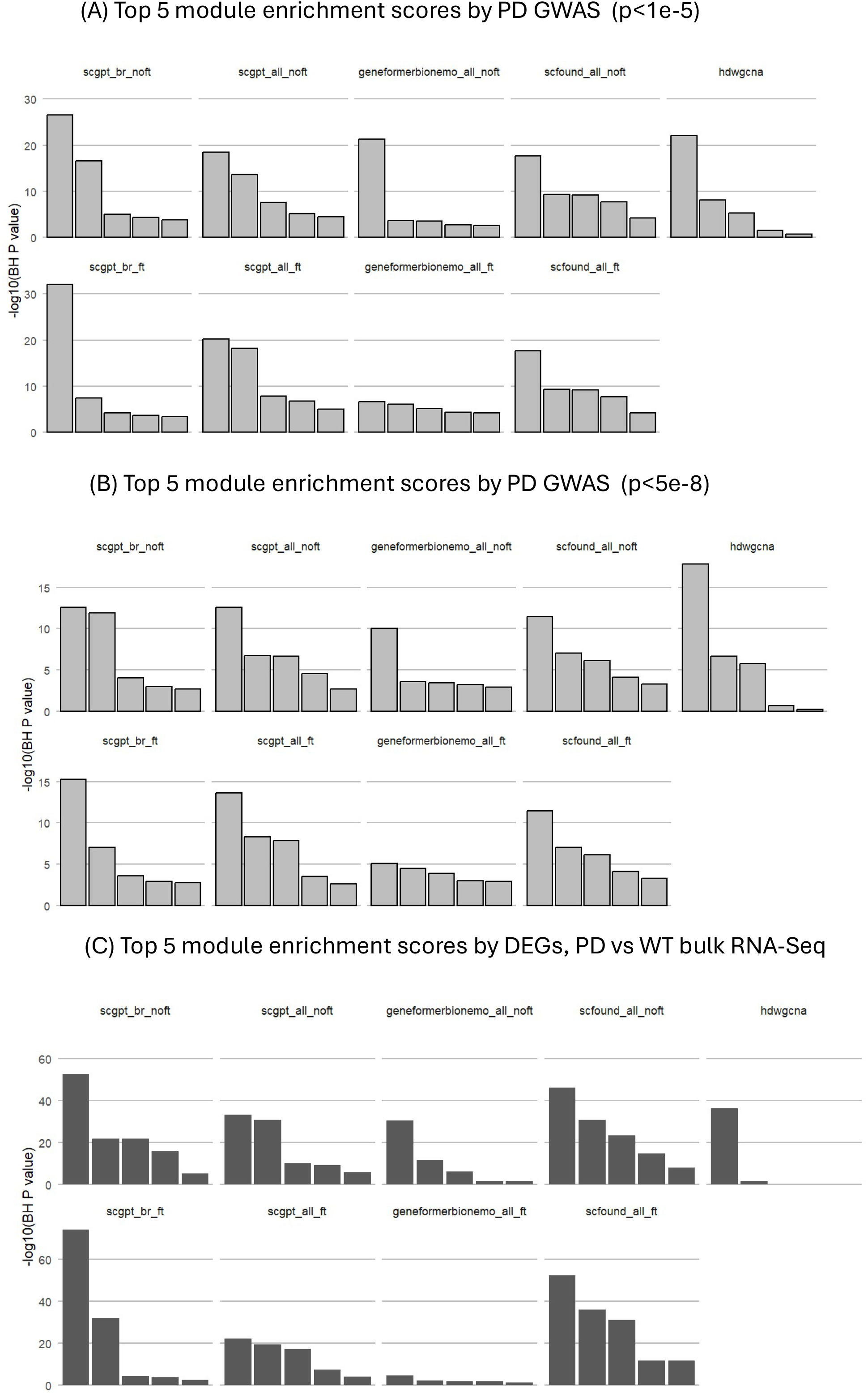
Benchmarking gene networks generated using foundation models and hdWGCNA. Enrichment analysis of gene modules using PD GWAS loci and PD organoid DEGs. Comparisons are shown for GWAS enrichment thresholds of P < 1×10⁻⁵ and P < 5×10⁻⁸ and enrichment of PD versus control organoid DEGs. The fine-tuned scGPT brain model shows the strongest enrichment for PD-associated signals. Model abbreviations indicate pre-trained (noft) and fine-tuned (ft) configurations.

**Supplementary Figure 2.**
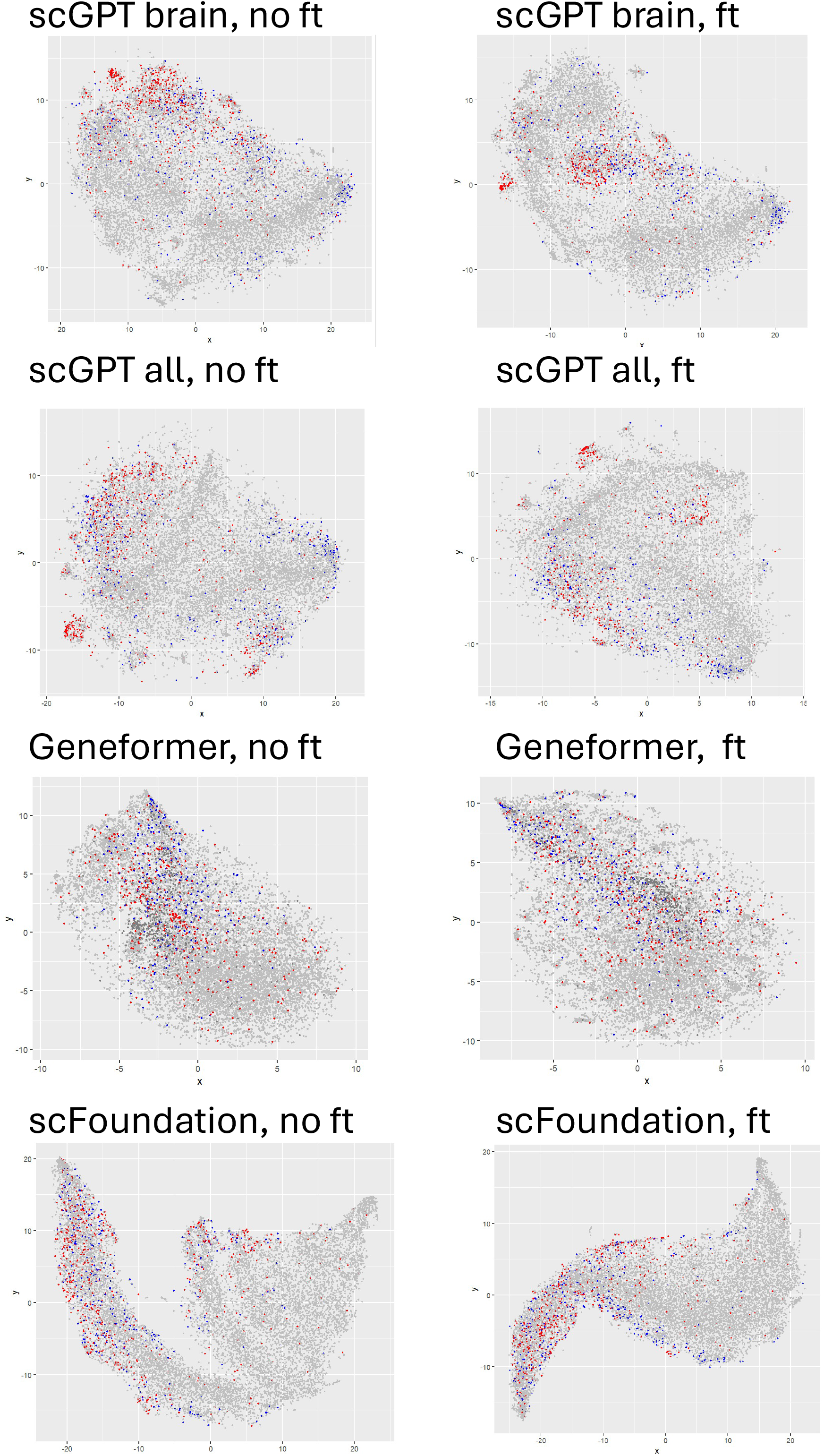
**Visualization of gene networks derived from pre-trained and fine-tuned foundation models**. t-SNE projections of gene embeddings for networks constructed using different foundation models. Differentially expressed genes from PD versus control organoid bulk RNA-seq are highlighted (red, upregulated; blue, downregulated).

**Supplementary Figure 3.**
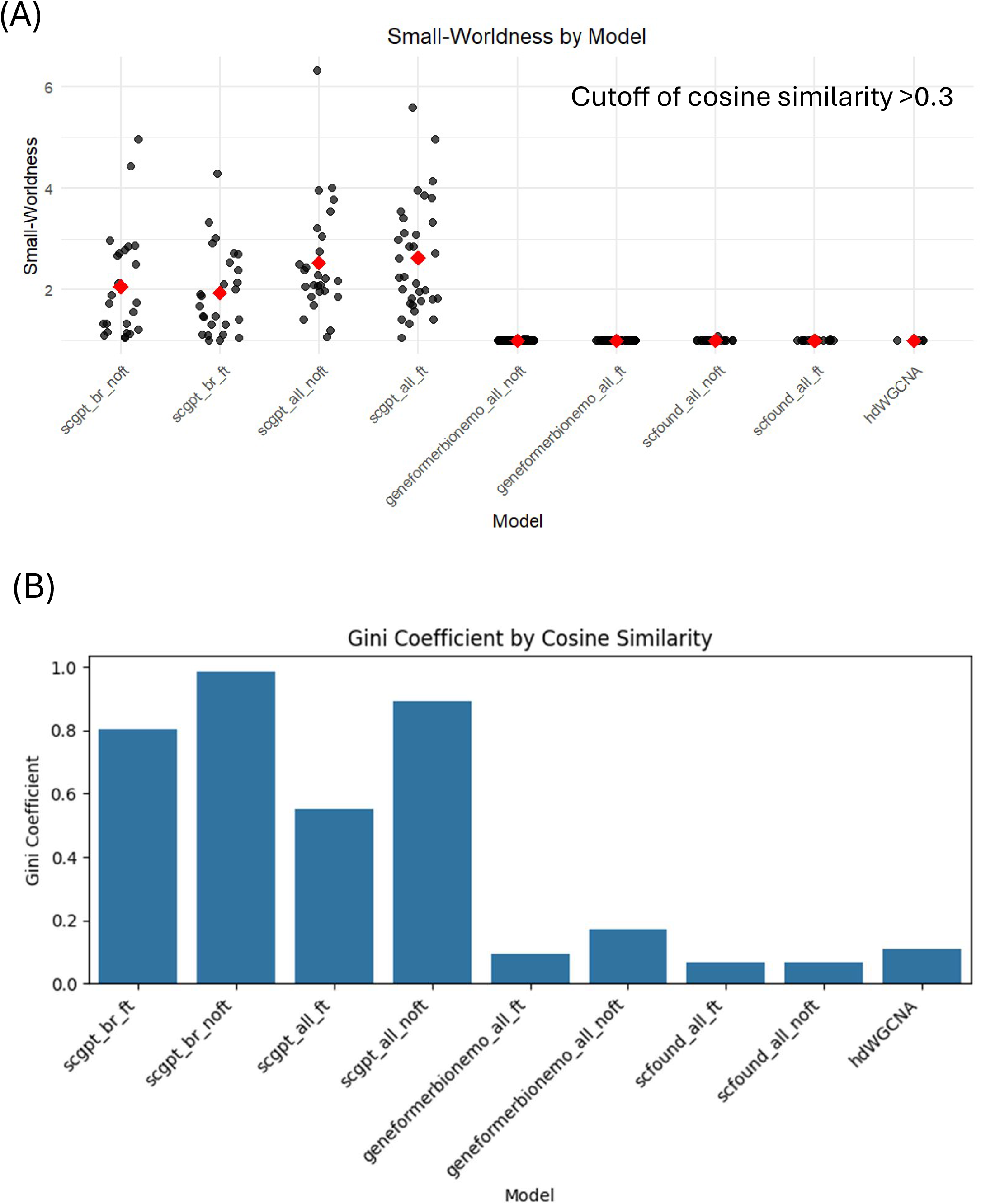
Topological benchmarking of foundation model–derived gene networks. (A) Small-worldness metric for gene networks generated using different foundation models and hdWGCN. (B) Gini coefficient quantifying node connectivity distribution across networks.

**Supplementary Figure 4.**
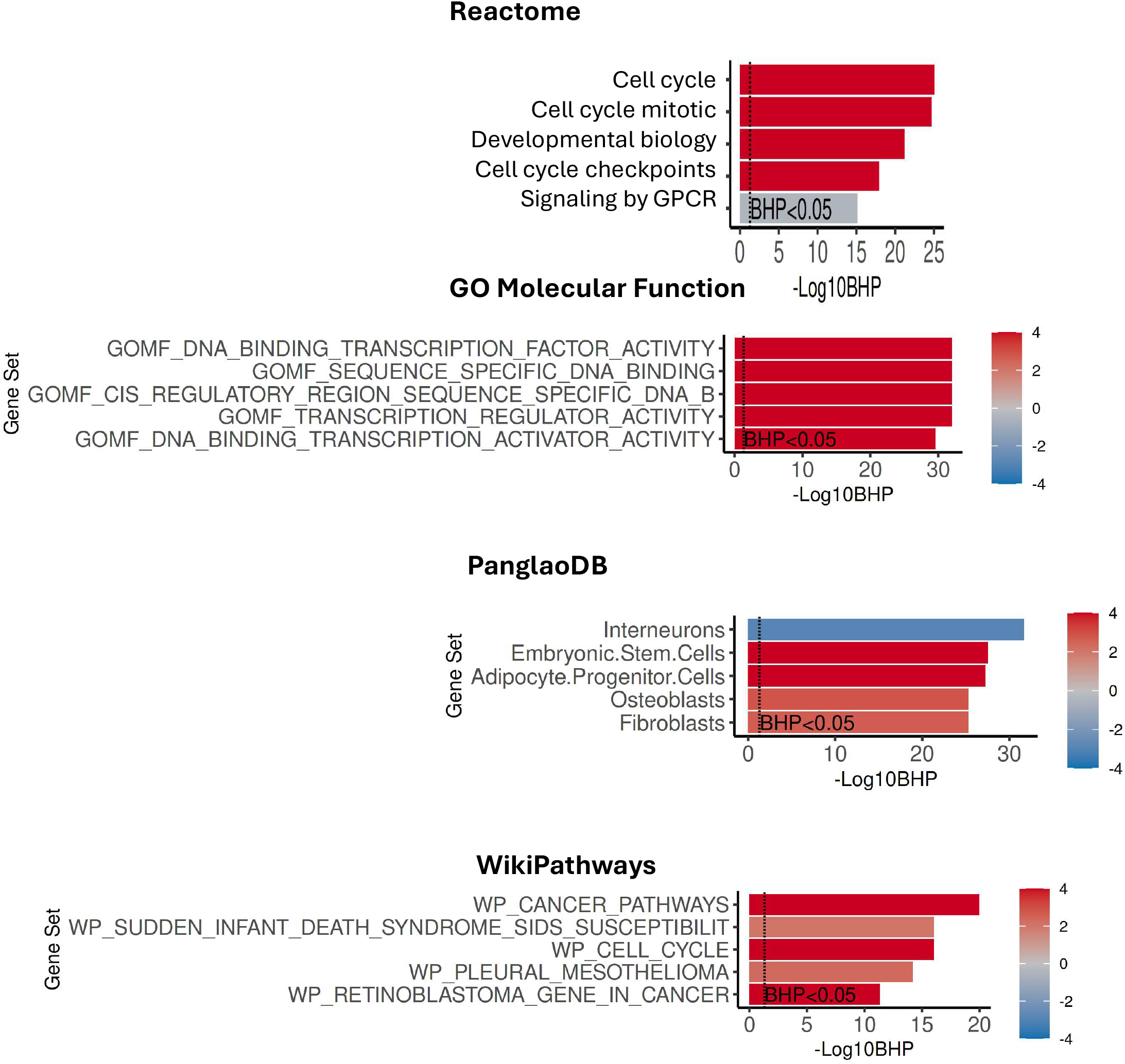
Functional enrichment of PD-associated genes. Gene set enrichment scores for Reactome, GO Molecular Function, PanglaoDB, and WikiPathways terms for DEGs between PD and control GBA1 organoids.

**Supplementary Figure 5.**
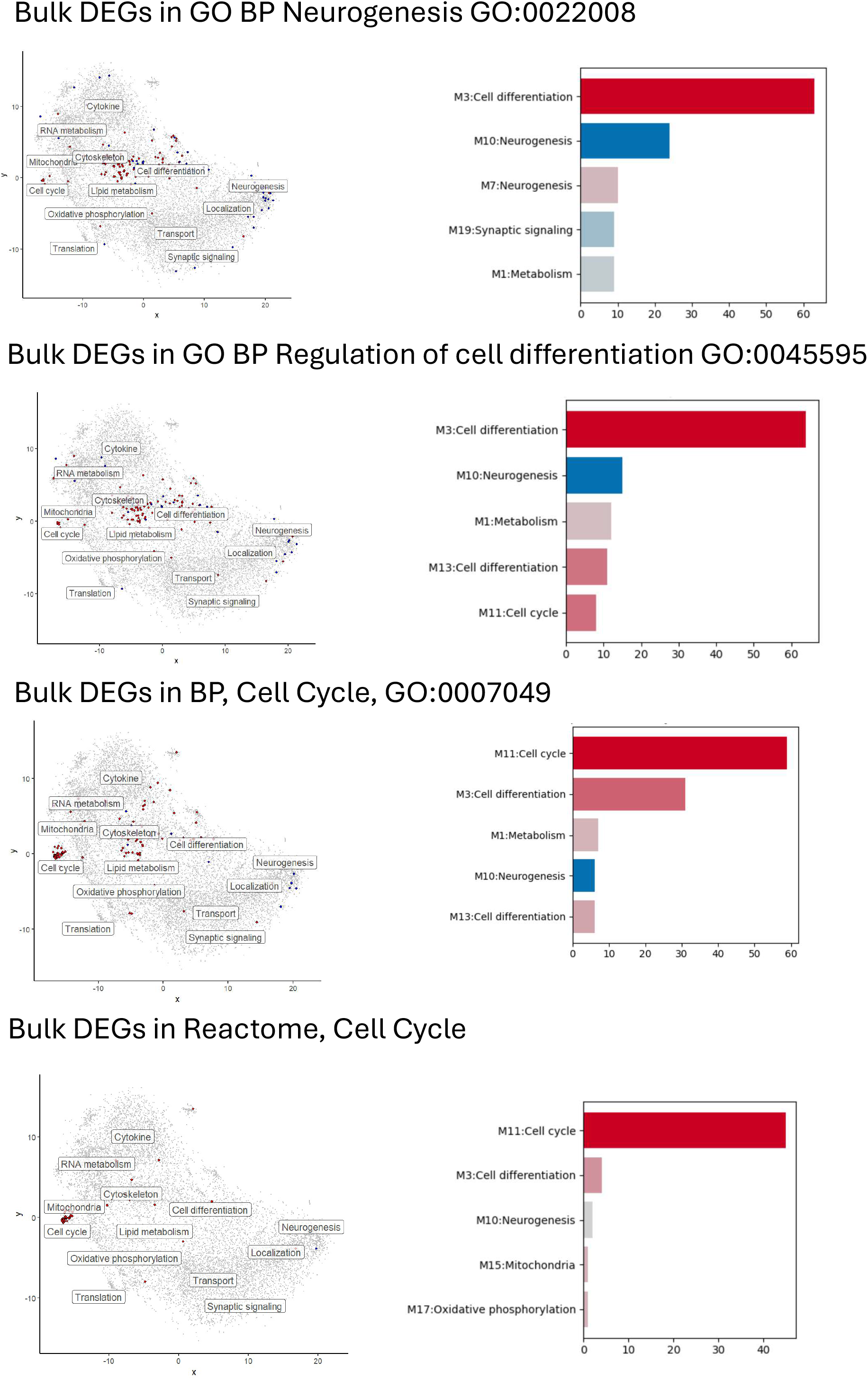
**Distribution of GO-associated genes across gene network modules**. Differentially expressed genes associated with selected GO terms mapped onto the gene network. Bar plots indicate the number of genes and enrichment Z-scores within each module.

**Supplementary Figure 6.**
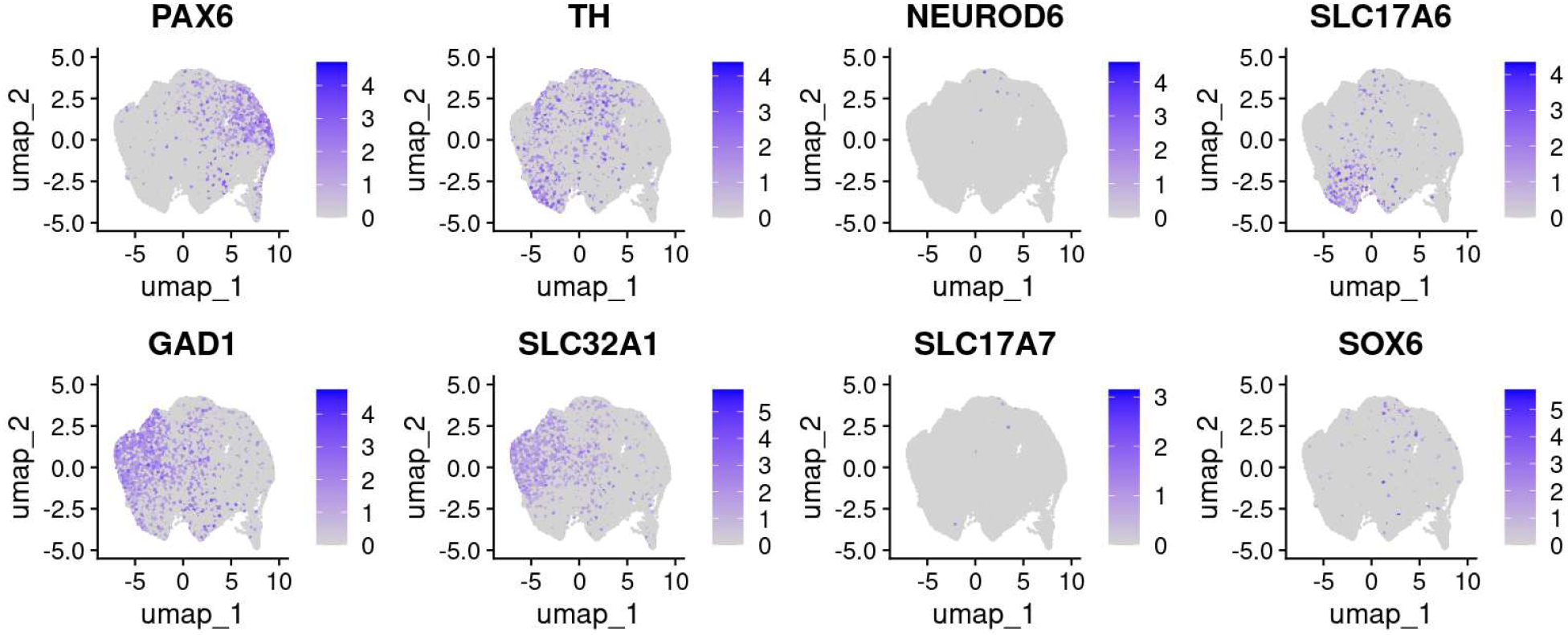
Cell-type marker expression across snRNA-seq clusters. UMAP visualization showing expression of marker genes for major cell types in PD and control midbrain organoids.

**Supplementary Figure 7.**
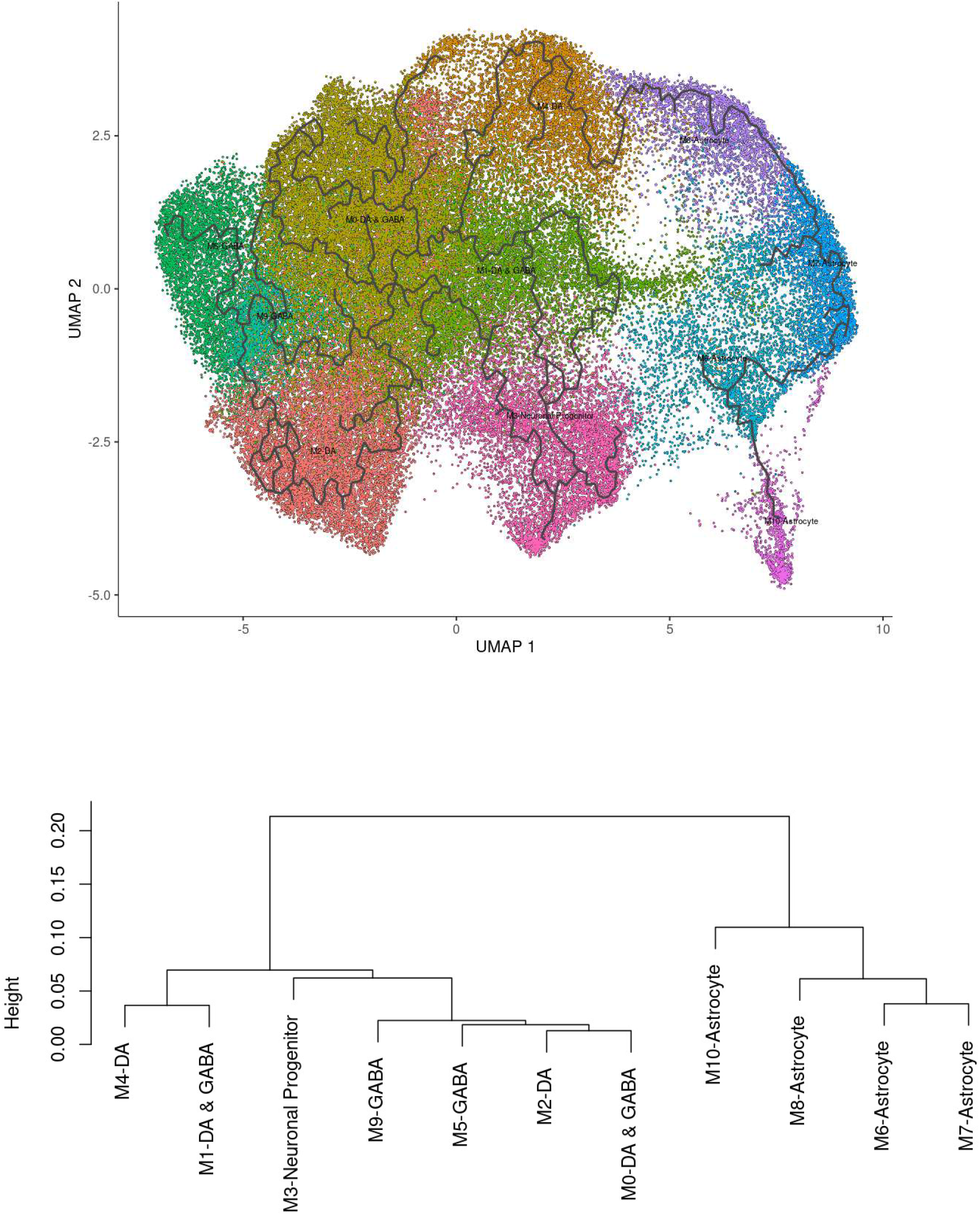
Cell trajectory analysis of midbrain organoid snRNA-seq data. Pseudotime trajectory analysis using Monocle 3 and hierarchical clustering of cell populations based on transcriptional similarity.

**Supplementary Figure 8.**
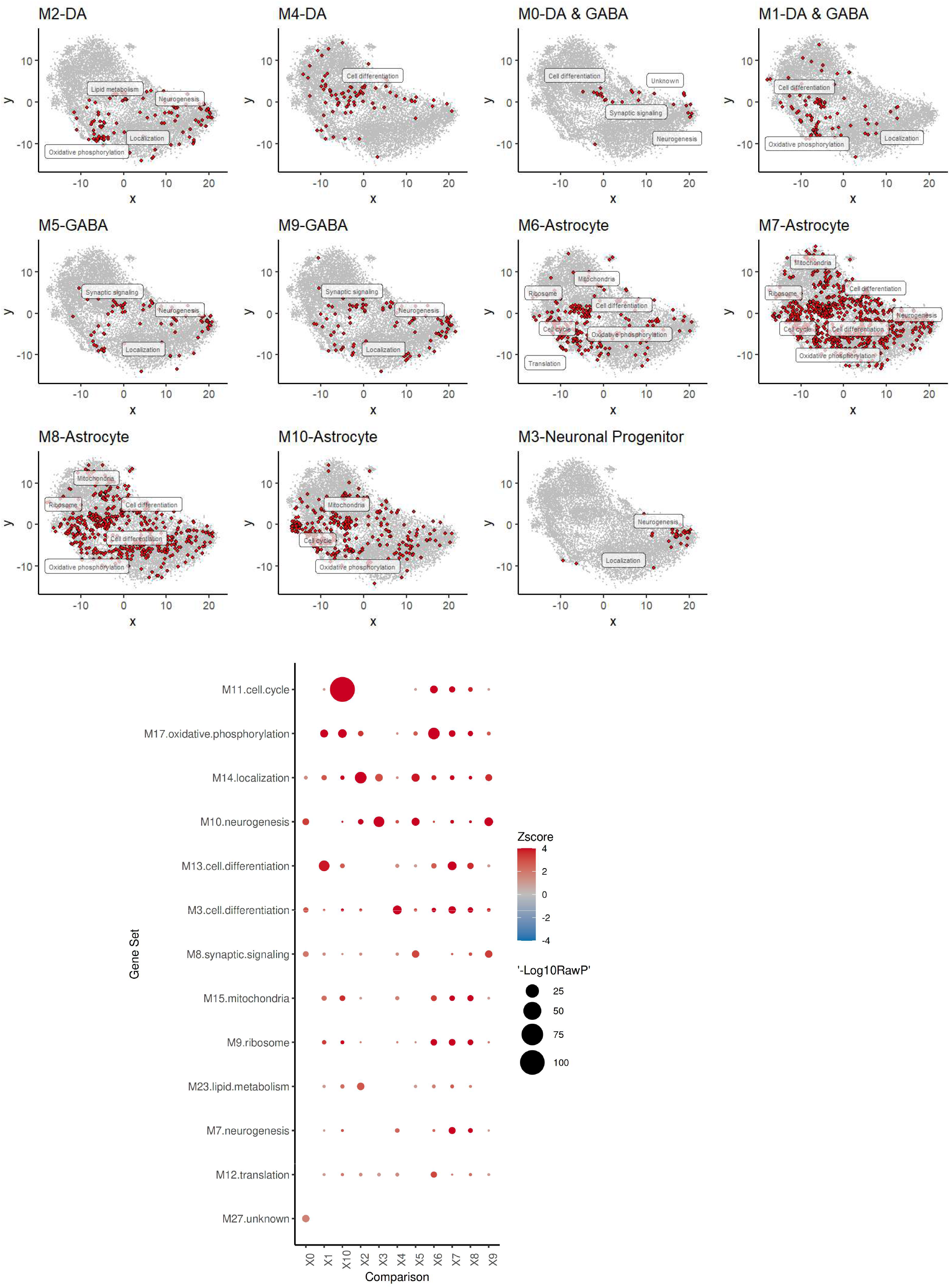
Cell-type marker mapping onto gene networks. Marker genes from different cell clusters mapped onto the foundation model–derived gene network.

**Supplementary Figure 9.**
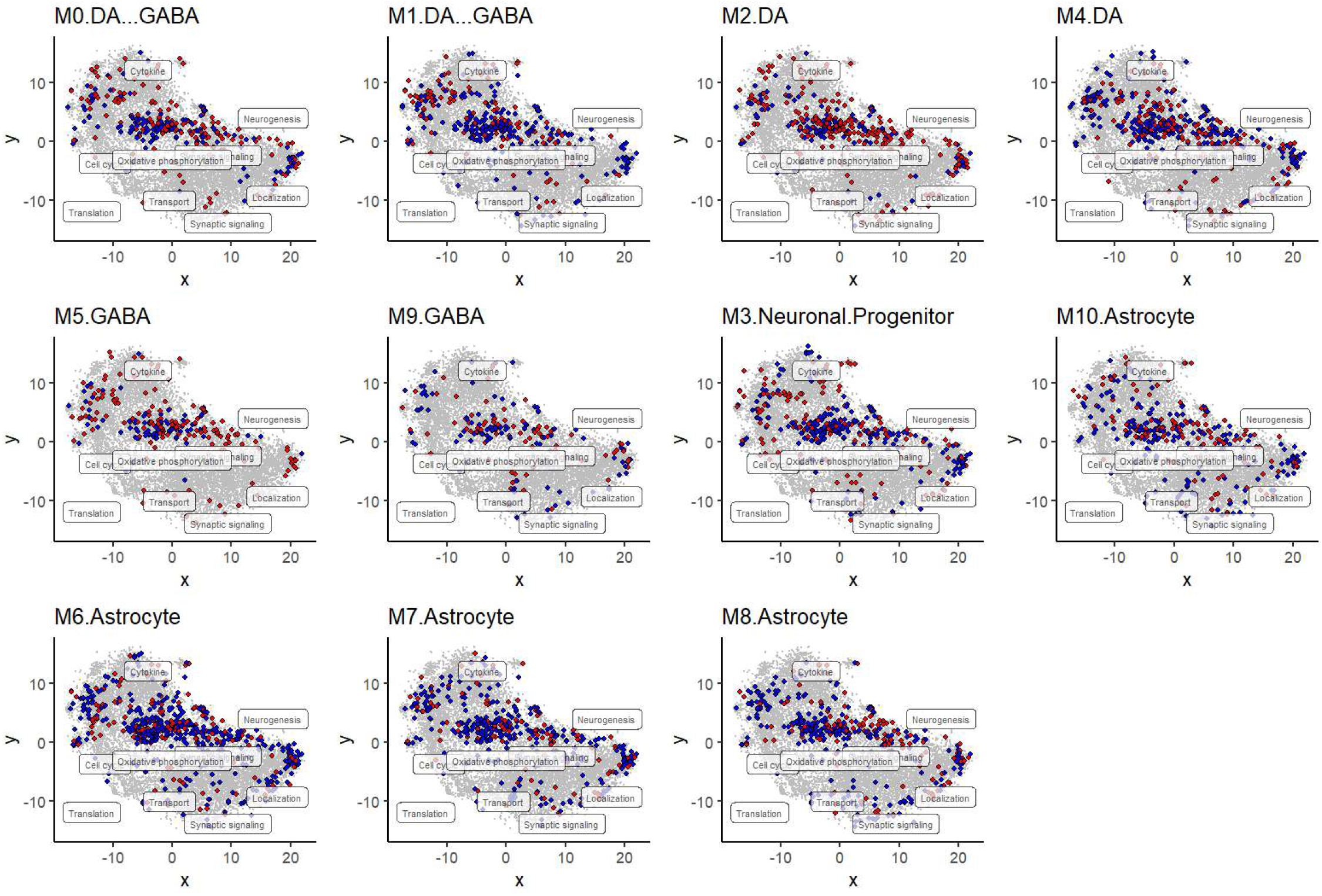
**Differential expression across cell clusters in PD organoids**. Differentially expressed genes between PD and control samples across cell clusters identified in snRNA-seq datasets.

**Supplementary Figure 10.**
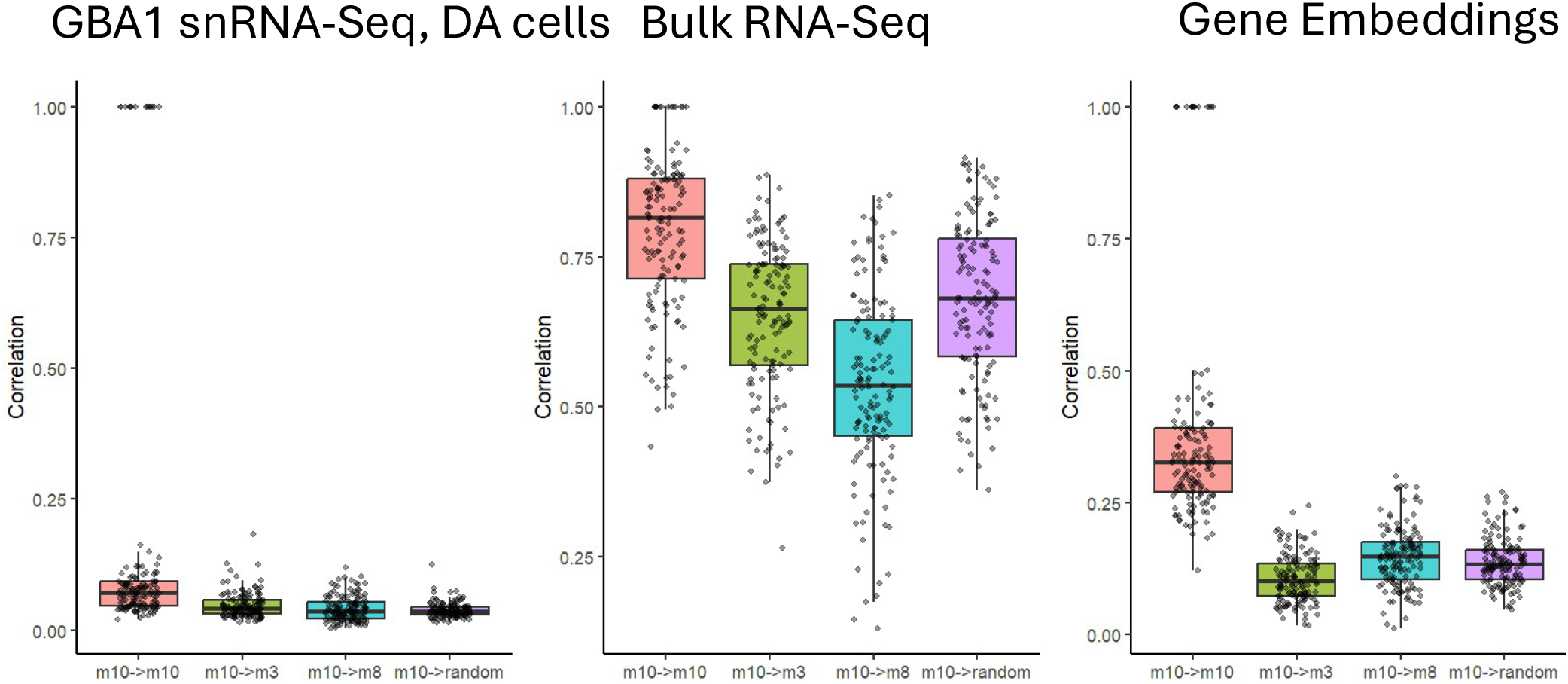
Cross-dataset correlation of neurogenesis module genes. Spearman correlation of DEGs from the M10 neurogenesis module in GBA1 organoids with DEGs in idiopathic PD brain datasets compared with genes from other modules or randomly selected genes.

**Supplementary Figure 11.**
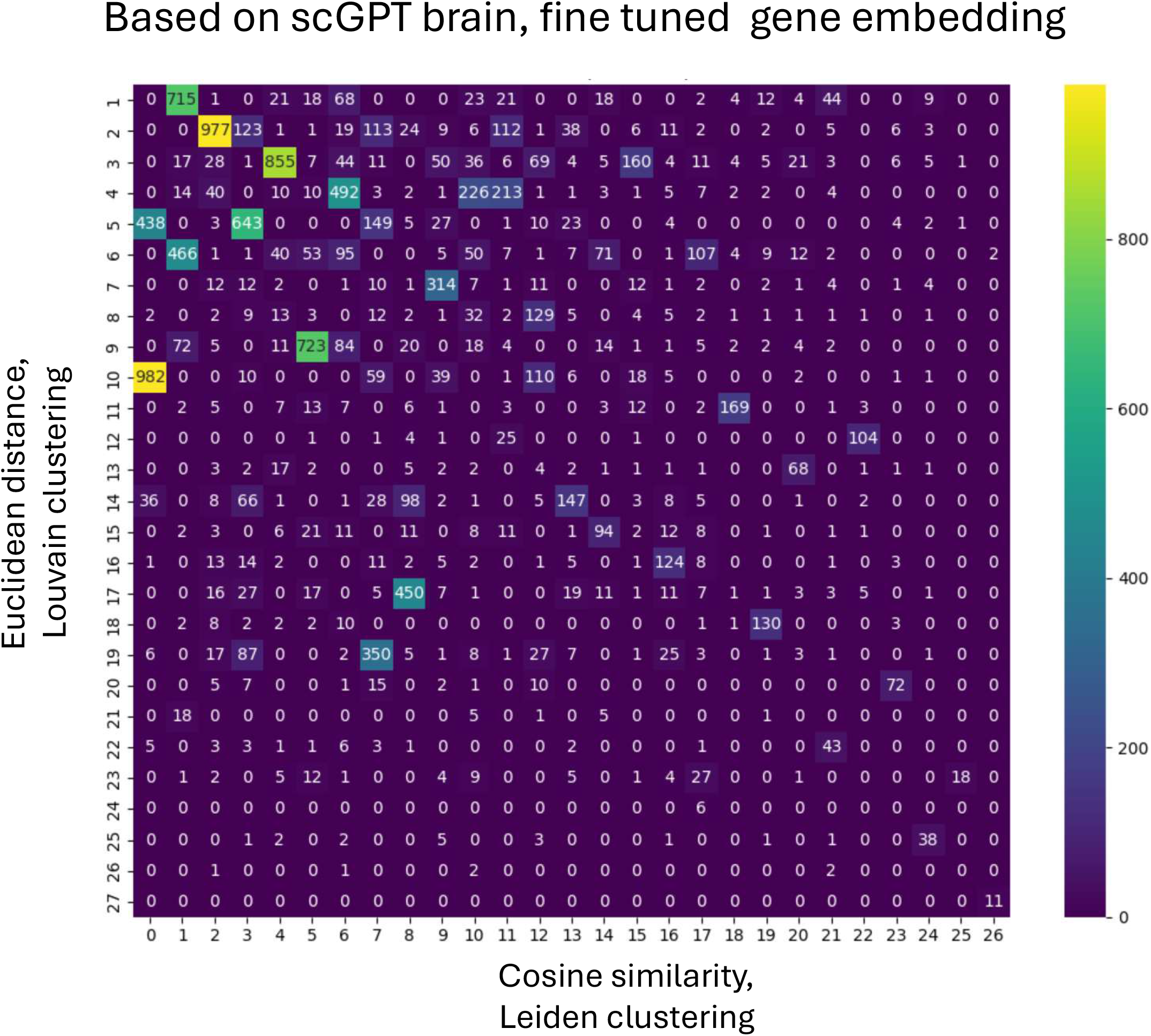
Robustness of gene module identification. Gene modules generated using alternative distance metrics and clustering methods show consistent clustering patterns.

## Methods

### Human iPSC culture and midbrain organoid differentiation

Midbrain organoids were generated from human induced pluripotent stem cells (iPSCs) following previously described protocols^16^. One iPSC line per donor was differentiated into neuroepithelial stem cells (NESCs), which were maintained in N2B27-based medium. NESCs were aggregated into spheroids to initiate organoid formation.

Organoids were patterned toward a midbrain fate using media containing small molecules and growth factors including ascorbic acid, CHIR-99021, purmorphamine, brain-derived neurotrophic factor (BDNF), and glial cell–derived neurotrophic factor (GDNF). Organoids were matured under these conditions to promote dopaminergic neuron differentiation. Organoids were collected at day 30 for downstream analyses.

### Single-nucleus RNA sequencing data processing

Single-nucleus RNA sequencing (snRNA-seq) count matrices were processed using the Seurat R package (version 4) ^27^. Gene expression counts were normalized using the standard Seurat workflow. Multiple datasets were integrated using canonical correlation analysis (CCA) implemented in the Seurat CCAIntegration method. Cells were clustered using the Leiden algorithm, and marker genes for each cluster were identified using the Wilcoxon rank-sum test. Cell-type annotation was performed by comparing cluster marker genes with previously reported markers for midbrain neuronal and glial populations30.

Differentially expressed genes (DEGs) between PD GBA1 and wild-type samples, PD LRRK2 and wild-type samples, or idiopathic PD and wild-type samples were identified using the Wilcoxon rank-sum test implemented in Seurat. For GBA1 and LRRK2 comparisons, DEGs were selected using a log2 fold change ≥0.5 or ≤−0.5 and a Benjamini–Hochberg (BH) adjusted P value <0.05. For idiopathic PD datasets, DEGs were selected using P <0.05.

### Bulk RNA sequencing analysis

Bulk RNA-seq count matrices were analyzed using the **DESeq2** R package^28^. Differential gene expression was assessed using the Wald test implemented in DESeq2. Genes were considered differentially expressed if they showed a log2 fold change ≥1 or ≤−1 with a BH-adjusted P value <0.05. Gene set enrichment analyses were performed using **Fisher’s exact test**. Z-scores were calculated as:

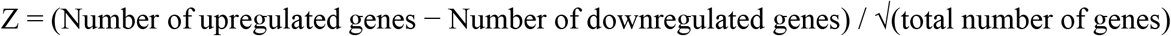

Positive Z-scores indicate enrichment of upregulated genes, whereas negative Z-scores indicate enrichment of downregulated genes.

### Fine-tuning of single-cell foundation models

Single-cell foundation models including scGPT9, Geneformer10, and scFoundation11 were fine-tuned using midbrain organoid snRNA-seq datasets containing more than 100,000 cells derived from seven familial PD patients with GBA1 mutations, one familial PD patient with an LRRK2 mutation, and three control donors.

The scGPT full model and the scGPT brain model were fine-tuned with a batch size of 16 for 5 epochs. The Geneformer 10M model from the NVIDIA BioNeMo framework^29^ was fine-tuned using default parameters in *train_geneformer*, and gene embeddings were extracted using *infer_geneformer*.

Due to memory constraints, the scFoundation model was fine-tuned using a subset of 30,000 cells and restricted to the final two layers, with a batch size of 16 for 10 epochs.

### Construction of foundation model–derived gene networks (scGENet)

Gene embeddings were extracted from fine-tuned foundation models and used to construct gene interaction networks. A k-nearest neighbor (kNN) graph was generated using cosine similarity between gene embeddings. Gene modules were identified using Louvain community detection.

Alternative distance metrics and clustering approaches implemented in the build_scGENet.py tutorial produced similar clustering patterns (Supplementary Fig. 11).

Gene modules were functionally annotated using enrichment analysis with Fisher’s exact test against public databases including Gene Ontology6, Reactome30, and PanglaoDB19.

For visualization, t-distributed stochastic neighbor embedding (t-SNE) was applied to reduce gene embedding dimensionality to two dimensions.

An interactive web application for exploring the gene network was developed using Python Dash and is available at:

https://www.brainstormtherapeutics.org/bstorganoidbrowser

### hdWGCNA co-expression network analysis

Gene co-expression networks were constructed using the hdWGCNA package^8^. Genes expressed in at least 5% of cells were retained for analysis.

Metacells were generated by aggregating transcriptionally similar cells using a k-nearest neighbor (kNN) approach. These meta cells were used to construct weighted gene co-expression networks with approximately scale-free topology following the hdWGCNA workflow.

### Network benchmarking and enrichment analysis

Gene modules derived from foundation model networks were evaluated by enrichment analysis using Gene Ontology biological processes, Parkinson’s disease GWAS loci, and differentially expressed genes from PD organoid bulk RNA-seq datasets.

Modules were ranked based on enrichment p-values from Fisher’s exact tests. Enrichment analyses were performed using PD-associated genes identified by GWAS with significance thresholds of P <1×10⁻⁵ and P <5×10⁻⁸.

### Network analysis of PD-associated genes

PD-associated genes were defined as genes differentially expressed between PD and wild-type samples. These genes were mapped onto the gene interaction network constructed using the fine-tuned scGPT brain foundation model.

Genes were visualized according to expression direction, with upregulated genes shown in red and downregulated genes shown in blue. Enrichment of DEGs within gene modules was assessed using Fisher’s exact test.

Network visualization and edge filtering were performed using Cytoscape31, with edges defined by cosine similarity ≥0.25.

### Cell-type deconvolution of bulk RNA-seq data

Cell-type proportions in bulk RNA-seq samples were estimated using the FARDEEP algorithm21.

Gene expression counts were normalized using counts per million (CPM). Reference expression profiles were derived from control samples in the snRNA-seq dataset. FARDEEP was run using default parameters with robust regression and adaptive outlier filtering to estimate cell-type composition.

### Cell trajectory analysis

Cell differentiation trajectories were inferred using Monocle 3^32^. UMAP embeddings generated using Seurat were imported into Monocle 3 to maintain the original low-dimensional representation.

The Seurat object was converted into a Monocle cell data set, and Monocle’s graph learning and pseudotime ordering algorithms were applied to infer differentiation trajectories and lineage relationships.

### Network topology analysis

Topological properties of gene interaction networks were evaluated using two metrics: small-worldness and the Gini coefficient.

Small-worldness (S) was calculated as:

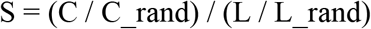

where C is the clustering coefficient of the observed network, L is the characteristic path length, and C_rand and L_rand represent the mean values from 100 degree-preserving randomized networks.

Gini coefficients were calculated to quantify inequality in node connectivity:

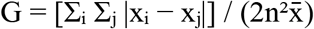

where xᵢ and xⱼ represent node degree values, x̄ is the mean node degree, and n is the number of nodes. Network analyses were performed using Python with the NetworkX and NumPy libraries.

### Statistics and reproducibility

Statistical analyses were performed using R and Python. Unless otherwise stated, statistical significance was assessed using two-sided tests. Multiple hypothesis testing corrections were performed using the Benjamini–Hochberg (BH) method.

## Data availability

Single-cell sequencing data from PD midbrain organoids are available through the NCBI Gene Expression Omnibus (GEO) under accession numbers GSE268784 (GBA1 organoids) and GSE133894 (LRRK2 organoids). Bulk RNA-seq data from PD midbrain organoids are available under accession numbers GSE287566 and GSE269316. Single-cell sequencing data from idiopathic PD midbrain samples are available under GSE157783. The ASAP Parkinson’s disease dataset was obtained from the ASAP CRN Cloud (cloud.parkinsonsroadmap.org).

## Code availability

Code for constructing scGENet foundation model–derived gene networks and reproducing the analyses is available at Github upon reviewer’s request.

## Computational resources

Model training and fine-tuning were performed using AWS EC2 instances equipped with NVIDIA T4 or L4 GPUs.

## Funding Statement

National Science Foundation SBIR Phase 1 grant number 2414877 to Robert T. Fremeau, Jr., Ph.D.

## Notes

### Competing Interest Statement

The authors have declared no competing interest.

